# Expectations and blind spots for structural variation detection from short-read alignment and long-read assembly

**DOI:** 10.1101/2020.07.03.168831

**Authors:** Xuefang Zhao, Ryan L. Collins, Wan-Ping Lee, Alexandra M. Weber, Yukyung Jun, Qihui Zhu, Ben Weisburd, Yongqing Huang, Peter A. Audano, Harold Wang, Mark Walker, Chelsea Lowther, Jack Fu, Human Genome Structural Variation Consortium, Mark B. Gerstein, Scott E. Devine, Tobias Marschall, Jan O. Korbel, Evan E. Eichler, Mark J. P. Chaisson, Charles Lee, Ryan E. Mills, Harrison Brand, Michael E. Talkowski

## Abstract

Virtually all genome sequencing efforts in national biobanks, complex and Mendelian disease programs, and emerging clinical diagnostic approaches utilize short-reads (srWGS), which present constraints for genome-wide discovery of structural variants (SVs). Alternative long-read single molecule technologies (lrWGS) offer significant advantages for genome assembly and SV detection, while these technologies are currently cost prohibitive for large-scale disease studies and clinical diagnostics (∼5-12X higher cost than comparable coverage srWGS). Moreover, only dozens of such genomes are currently publicly accessible by comparison to millions of srWGS genomes that have been commissioned for international initiatives. Given this ubiquitous reliance on srWGS in human genetics and genomics, we sought to characterize and quantify the properties of SVs accessible to both srWGS and lrWGS to establish benchmarks and expectations in ongoing medical and population genetic studies, and to project the added value of SVs uniquely accessible to each technology. In analyses of three trios with matched srWGS and lrWGS from the Human Genome Structural Variation Consortium (HGSVC), srWGS captured ∼11,000 SVs per genome using reference-based algorithms, while haplotype-resolved assembly from lrWGS identified ∼25,000 SVs per genome. Detection power and precision for SV discovery varied dramatically by genomic context and variant class: 9.7% of the current GRCh38 reference is defined by segmental duplications (SD) and simple repeats (SR), yet 91.4% of deletions that were specifically discovered by lrWGS localized to these regions. Across the remaining 90.3% of the human reference, we observed extremely high concordance (93.8%) for deletions discovered by srWGS and lrWGS after error correction using the raw lrWGS reads. Conversely, lrWGS was superior for detection of insertions across all genomic contexts. Given that the non-SD/SR sequences span 90.3% of the GRCh38 reference, and encompass 95.9% of coding exons in currently annotated disease associated genes, improved sensitivity from lrWGS to discover novel and interpretable pathogenic deletions not already accessible to srWGS is likely to be incremental. However, these analyses highlight the added value of assembly-based lrWGS to create new catalogues of functional insertions and transposable elements, as well as disease associated repeat expansions in genomic regions previously recalcitrant to routine assessment.

## Main Text

The field of genomics has seen remarkable advances in the accuracy and efficiency of massively parallel sequencing-by-synthesis technology that generates pairs of short reads from the ends of small 400-800 base pair (bp) fragments (referred to herein as short-read WGS [srWGS]). This technical leap, and derivative approaches such as targeted exome capture sequencing (WES), have catalyzed a deluge of gene discoveries for rare diseases and insights into population genetics and genome biology. Correspondingly, srWGS has been adopted by all major human disease and biobank sequencing initiatives, including the NHGRI Centers for Common Disease Genomics (CCDG)^1^ and Centers for Mendelian Genetics (CMG),^2^ the Deciphering Developmental Disorders (DDD) project,^3^ the Trans-Omics for Precision Medicine (TOPMed),^4^ the All of Us Research Program,^5^ the NICHD Gabriella Miller Kids First (GMKF) initiative, the UK BioBank,^6^ and Genomics England,^7^ to name just a few. As such, a critical step for the field is to establish uniform methods for srWGS data processing and rational benchmarking standards to set expectations for variant detection.

The technical processes of genome alignment and single nucleotide variant (SNV) detection have been an intensive focus of genomics since the inception of the 1000 Genomes Project,^8–10^ and more recently updated for cross-institute functional equivalence as part of the NHGRI Genome Sequencing Program.^11^ However, no standardized methods have been adopted for structural variants (SVs), defined as genomic alterations greater than 50 bp in size, and consequently no gold-standard benchmarking approaches exist for SV discovery. This lack of uniformity has introduced a barrier to the establishment of reliable estimates of the SV counts and characteristics per genome that are comparable to those established for short variants. Not surprisingly, as shown in Figure 1A these estimates have varied considerably across studies. The initial discovery effort from the 1000 Genomes Project^12,13^ revealed that a diverse landscape of SVs could be captured from srWGS with just 4-7X coverage (3,422 SVs per genome), and more recent population genetic and human disease studies using deeper (30X or higher) srWGS and diverse methods have varied in estimates of SVs that can be captured using srWGS from 401 – 10,884 per genome, with the highest end of this range generated from the Human Genome Structural Variation Consortium (HGSVC; Figure 1A).^1,13–18^

**Figure 1.**
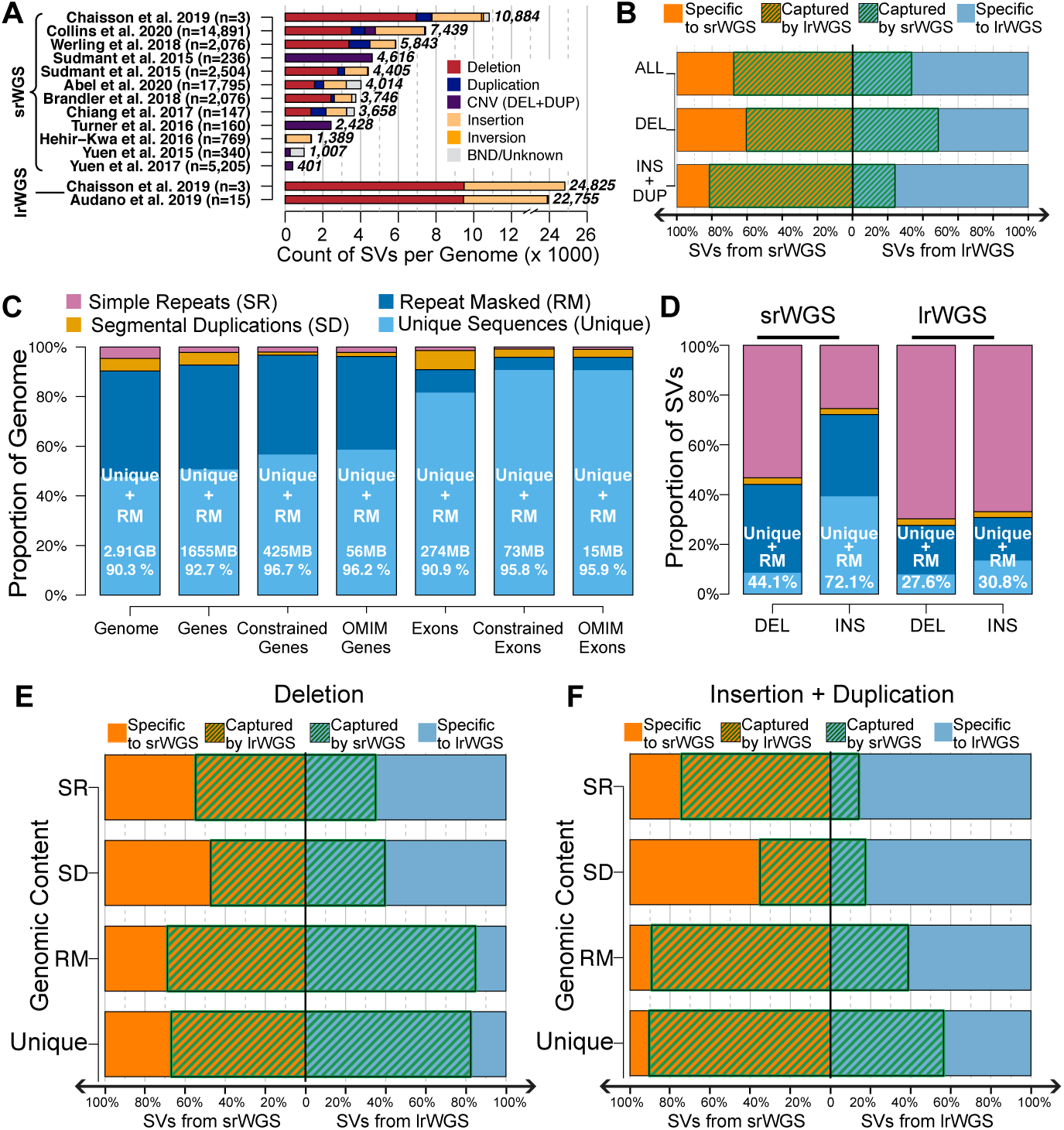
Comparison of SV callsets from srWGS and lrWGS. (A) The substantially increased yield of lrWGS in SV detection is displayed from the HGSVC (Chaisson et al 2019)^14^ and the largest Pacific Biosciences (PacBio) lrWGS study published to date (Audano et al 2019)^22^ by comparison to contemporary srWGS studies. As shown, there is wide variability of SV detection across srWGS studies to date that report SVs detected per individual in more than 100 genomes. Parentheses next to each study label indicate the number of genomes analyzed, and bold numbers next to each bar represent the number of SVs per genome reported by each study. (B) Overlap of SVs from the HGSVC srWGS and lrWGS callsets across children of the three trio families, partitioned by SV class. (C) Distribution of repetitive sequences across the genome, genes, and exons. Constrained refers to genes and exons with pLI > 0.9,^36^ and OMIM Genes include a curated list of autosomal dominant genes that were defined in both Berg et al.^43^ and Blekhman et al.^44^ GB = gigabase, MB = megabase. Percentage listed within each bar is the fraction of each group composed of Unique + RM sequences. (D) Distribution of SVs from srWGS and lrWGS split by repetitive sequence context. Formatting conventions are the same as panel C. (E-F) Concordance of (E) deletions and (F) insertions and duplications between srWGS and lrWGS split by repetitive sequence context.

Emerging long-read WGS (lrWGS) technologies, which involve sequencing thousands to millions of contiguous nucleotides from a single strand of DNA, are better suited for SV discovery than srWGS. The most widely tested lrWGS technologies include single-molecule real-time (SMRT) sequencing from Pacific Biosciences (PacBio) and sequencing by ionic current through a nanopore channel (Oxford Nanopore Technologies [ONT]). A key advantage of lrWGS is the abundance of reads that span entire SVs, allowing for direct observation rather than detection by inference as required for srWGS. These unique properties of lrWGS are beginning to revolutionize *de novo* assembly approaches,^19,20^ with methods already maturing for telomere-to-telomere assembly of individual human chromosomes.^21,22^ The most recent lrWGS analyses have at least doubled the number of SVs able to be captured in each genome to ∼25,000 as compared to srWGS^14,22^ (Figure 1A). The impact of these studies has exceeded the sheer volume of variants detected: assembly-based long-read analyses have opened access to variants in the genome that have been traditionally refractory to delineation by short read sequencing or interpretation in disease association studies, such as repeat expansions and alterations within repetitive segmental duplications and centromeres.^23^ Unfortunately, the current cost of lrWGS is a significant premium over srWGS, depending on the technology used. By example, the current cost for generation of PacBio lrWGS over srWGS for equivalent coverage at leading academic platforms from the HGSVC ranges from 5.9-fold increase for continuous long read (CLR) technology to 12-fold increase for circular consensus sequencing (CCS) HiFi technology. Moreover, the low throughput of modern lrWGS platforms renders them impractical for adoption in most large-scale population studies. The largest published assembly-based PacBio study has analyzed just 15 genomes,^22^ while a recent study from Iceland analyzed 1,817 ONT genomes,^24^ by comparison to millions of genomes that have already been sequenced or commissioned using srWGS. Given this predominance of srWGS in the current landscape of genomics research, we present here a series of analyses from the HGSVC to: (i) define and quantify the limitations of SV detection from srWGS; (ii) benchmark expectations for the number and class of variants that can be reliably detected from srWGS; (iii) predict the genomic features that drive false positive and false negative discoveries for each technology; and (iv) establish the scientific and clinical advances offered by state-of-the-art lrWGS assembly as a complementary approach to srWGS.

In this study, we performed a detailed comparison of SV detection from alignment-based srWGS and assembly-based lrWGS methods on matched samples. In the HGSVC, we recently generated SV callsets from srWGS and lrWGS of three parent-child trios from the 1000 Genomes Project.^14^ For srWGS, this initial HGSVC study applied a highly sensitive ensemble approach, involving 13 SV detection algorithms (Supplemental Methods), and discovered 10,884 SVs per genome. The emphasis on sensitivity suggests that ∼11,000 SVs per genome likely reflects an upper bound on the total number of SVs that can be captured from srWGS with alignment-based algorithms applied by the HGSVC, as demonstrated in Figure 1A by comparison to other contemporary studies. However, this sensitivity came at the significant cost of specificity, with 685 *de novo* SVs observed per genome, or >1,000-fold more than expected from srWGS based on family studies, population genetic estimators, and molecular validation, therefore representing many variant predictions that are likely false positives.^15,16,25^ The lrWGS-derived SV callset combined whole genome phasing with two state-of-the-art genome assembly approaches (Phase-SV and MS-PAC^19,20,26^) and was supplemented by additional technologies (HiC and StrandSeq, see Chaisson et al.^14^). These methods discovered an average of 24,825 haplotype-resolved SVs per genome, or over two-fold more than the most sensitive srWGS approaches. Surprisingly, although the srWGS and lrWGS callsets were generated on identical samples, only a limited subset of SVs (66.7% of srWGS and 33.5% of lrWGS) overlapped between technologies. Moreover, the mutational class of SVs dramatically impacted concordance: 60.5% of srWGS and 48.7% of lrWGS deletions demonstrated overlap as compared to 81.5% of srWGS and 24.1% of lrWGS insertions (Figure 1B).

We sought to define and quantify the factors contributing to the poor concordance between SVs derived from each technology on matched samples, as these factors might be used to improve SV discovery, filtering, and prioritization in medical and population genetic initiatives. We first explored the role of genomic features such as repetitive sequences that are enriched for SVs due to repeat-mediated mechanisms,^22,27,28^ as short-read alignment has well-documented limitations within these genomic regions.^29,30^ We annotated all SVs with sequence context based on RepeatMasker^31^ and segmental duplication^32,33^ tracks from the UCSC genome browser.^34,35^ For simplicity, we consolidated all repetitive sequence annotations into three categories: segmental duplication (SD; 5.1% of the genome), simple repeat (SR; 4.6%), and referred to all other repetitive sequence not overlapping SD/SR elements as ‘repeat masked’ (RM; 42.9%). The remaining 47.4% of the genome not overlapping any of these repeat categories was labeled as ‘Unique’ sequence, which is a term used for simplicity here but these regions are not completely devoid of some duplicated sequences. The Unique and RM categories collectively encompass 90.3% of the annotated human reference sequence, 90.9% of all currently annotated protein-coding sequence, 95.8% of all currently annotated coding sequence from evolutionarily constrained genes, and 95.9% of genes currently associated with human disease from the Online Mendelian Inheritance in Man (OMIM; Figure 1C).^36–39^

As expected, the distribution of SVs was non-uniform and varied by sequence context for each technology (Figure 1D). Most prominently, the enrichment of SV breakpoints in highly repetitive genomic sequences (SD/SR regions) was dramatic and their distribution differed significantly between technologies: despite representing just 9.7% of the reference genome, SD/SR annotated sequences contained at least one breakpoint from 49.8% of all SVs from srWGS and 70.4% of all SVs from lrWGS (P < 2.2e-16 for both technologies, chi-square test, Table S2, see Supplemental Methods for details). This enrichment of SVs in repetitive sequence was also strongly correlated with concordance between srWGS and lrWGS, as SVs located in repetitive SD/SR sequences displayed 57.0% concordance among srWGS variants and 22.5% in lrWGS variants, whereas those ratios improved considerably in less repetitive sequences (Unique + RM) to 76.5% in srWGS and 59.9% in lrWGS (Figure 1E-F).

While the divergent distributions and diminished concordance of SV detection by technology aligned with expectations for SD/SR regions, the paucity of overlap between technologies in Unique + RM regions was unexpected as breakpoints localized to these regions should not suffer from the technical confounds that profoundly impact SV discovery in highly repetitive sequences. We next sought to decouple and quantify the discordance driven by underlying biological features of the genome from technical noise driven by false positive SVs present in the underlying HGSVC callsets, which were optimized for sensitivity as described above. We also reasoned that determining the covariates that have the greatest influence on false positive calls would be of high value. To accomplish this, we developed an *in silico* SV assessment procedure to improve the precision of srWGS and lrWGS callsets in non-repetitive regions. This procedure re-evaluated the following three pieces of orthogonal information from both lrWGS and srWGS for each SV: (1) supporting evidence from an algorithm that surveys the raw lrWGS reads for the presence of an SV (VaPoR;^40^ Figure 2A); (2) copy states based on srWGS normalized read depth (RD) within SVs (Figure 2B, S1); (3) discordant paired-end (PE) and split reads (SR) at the breakpoint of each predicted SV (Figure 2C-D, S2, Table S1). We considered the SVs with one or more modes of supporting evidence as “high confidence” and explored their overlap based on repeat context for SV calls from different technologies (see Supplemental Methods for further details).

**Figure 2.**
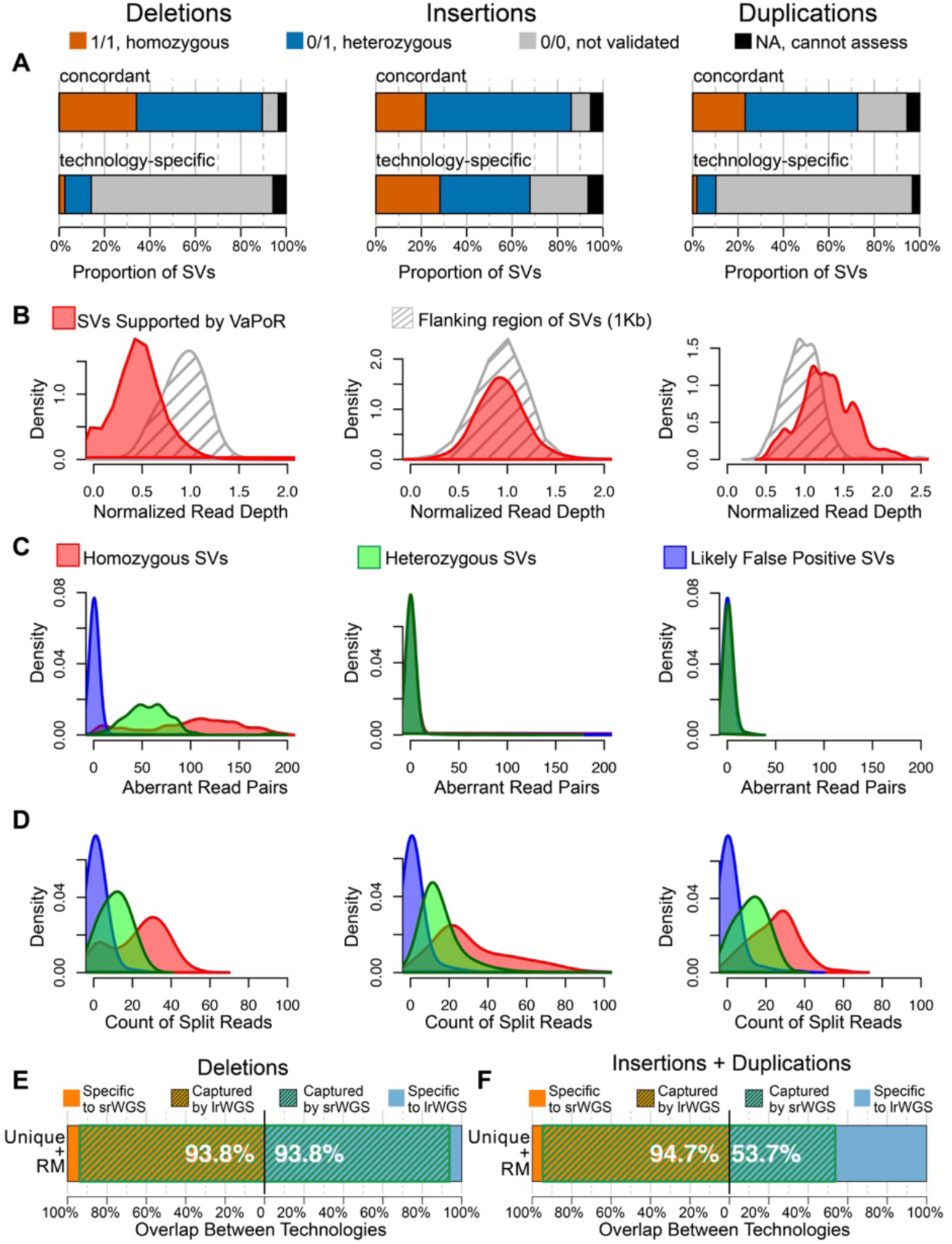
Error correction methods for SVs in Unique + RM region and the updated concordance. (A) In silico evaluation results from VaPoR on deletions (left), insertions (middle) and duplications (right). Deletions and insertions were reported in both srWGS and lrWGS callsets; duplications were only reported in the srWGS callset. (B) Distribution of normalized read depth of srWGS across deletions (left), insertions (middle) and duplications (right) that were supported by VaPoR (red), and the 1Kb genomic regions that flank each SV (grey). (C-D) Distribution of (C) aberrant srWGS read pairs and (D) split reads around deletions (left), insertions (middle) and duplications (right) that were either homozygous (red), heterozygous (green) or false positives (blue). The homozygous, heterozygous and likely false positive SV sets were selected using the criteria described in supplemental methods. (E-F) Concordance of (E) deletions and (F) insertions and duplications in Unique + RM sequences that were supported by the in silico SV refinement procedure. Percentages represent the fraction of total variants shared between srWGS and lrWGS.

We initially applied this *in silico* SV refinement procedure to deletions, which represent the most interpretable class of SVs for genomics applications (Figure S3). As expected, the *in silico* confirmation rate—*i*.*e*., the proportion of SVs supported by one or more of the evidence classes described above—was high (93.5%) for deletions concordant between technologies in Unique + RM regions, compared to just 13.5% and 33.1% for those that were only discovered by a single technology for srWGS or lrWGS, respectively (Figure S4). After restricting to high confidence deletions with supportive information, just 6.2% of the deletions in Unique + RM regions were specific to either srWGS or lrWGS (Figure 2E). Although we cannot rule out explanations such as somatic SVs or sub-clonal mutations arising in cell culture, these results imply that the most of the discordance reported between srWGS and lrWGS for deletion discovery in the 90.3% of the genome not encompassed by SD/SR sequence was likely technical and driven by false positive SV calls that can be pruned by *post hoc* heuristic filtering.

In contrast to this strong concordance between srWGS and lrWGS observed for deletions, nearly half (46.3%) of high confidence lrWGS insertions in Unique + RM regions had no matching SV call from srWGS, while the majority (94.7%) of srWGS insertions and duplications were captured by lrWGS SV calls (Figure 2F, S5). To further investigate the properties of insertions specifically captured by lrWGS in Unique + RM sequences, we aligned the assembled sequences of high-confidence insertions against a catalog of known repeat elements.^31^ Most of these insertions aligned to specific types of repeat elements (61.8%, N = 2,485 / genome), such as short and long interspersed nuclear elements (SINEs, N = 1,494 / genome; LINEs, N = 312 / genome) and long terminal repeat (LTR, N = 139 / genome) retrotransposons (Figure 3A,D). Yet another 19.0% of the insertions exhibited partial alignments to multiple different repeat types (Figure 3A, C). Notably, most (70.1%) of the lrWGS insertions that were shared by srWGS aligned to a specific type of repeat element, whereas nearly one-third (31.7%) of the insertions specifically discovered by lrWGS were partially aligned to multiple different repeats types (Figure 3B, C), indicating that the complexity of chimeric repeat structures is a major determinant of srWGS sensitivity for insertion SVs, as has been previously demonstrated in certain classes of nested insertions.^41^ We further observed high variability in the current capabilities of srWGS detection depending on the type of transposable element insertions when comparing with lrWGS as 74.4%, 44.2% and 50.7% of lrWGS insertions were discovered by srWGS for SINEs, LINEs and LTRs, respectively (Figure 3D). Intriguingly, 95.8% of the high confidence lrWGS insertions in Unique + RM regions that did not overlap an srWGS insertion nevertheless had some detectable support in the raw srWGS data, indicating that continued development of detection algorithms could improve sensitivity for these missed insertion SVs (Figure 3E). Taken together, these analyses indicate that lrWGS and assembly-based approaches provide substantial improvements over srWGS for insertion discovery, particularly for those events with complex repeat structures.

**Figure 3.**
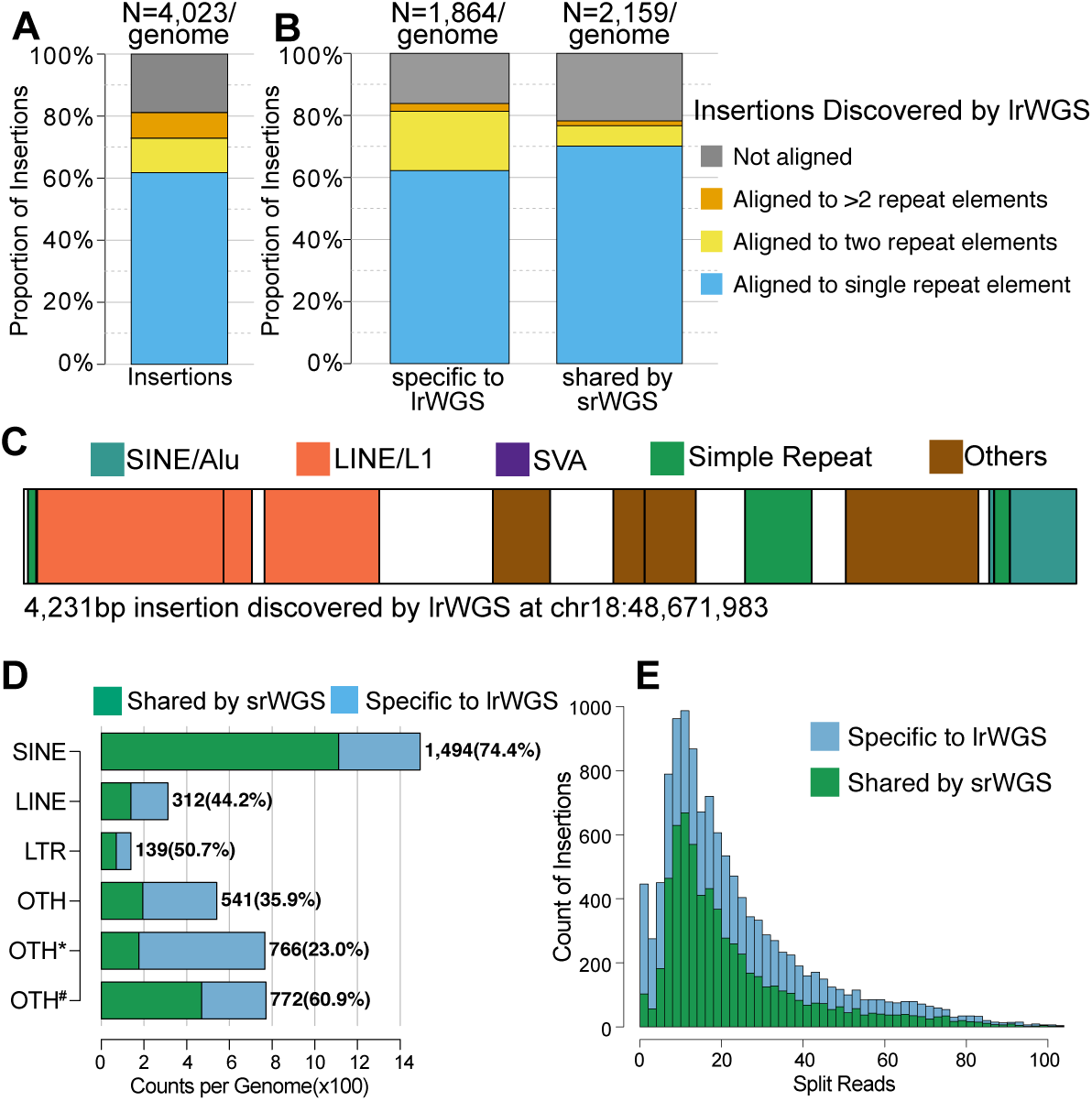
Alignment of assembled lrWGS insertion sequences against known repeat elements. (A) Count of lrWGS insertions in Unique + RM sequences per genome by alignment of inserted sequence to known repeat elements. Number on top of bar represents the averaged count of high confidence insertions in Unique + RM sequences per genome. (B) Count of lrWGS insertions that are specifically discovered by lrWGS and shared by srWGS, by alignment of inserted sequences to known repeat elements. Formatting conventions used are the same as panel A. (C) Example of an insertion SV assembled by lrWGS, annotated with sequences that align to known repeat element classes. (D) Counts of lrWGS insertions in Unique + RM sequences per genome by the class of inserted sequence and the proportion of variants that overlap with srWGS. “OTH*” represents insertions aligned to multiple known repeat elements, as the example shown in panel (B). “OTH#” represents insertions not aligned to any repeat elements. Number in parentheses represents the proportion of insertions that overlap with srWGS. (E) Count of split reads around the lrWGS high confidence insertions displayed in the histogram.

Finally, we examined SVs in highly repetitive SD/SR regions using the same *in silico* evaluation framework (Figure S6A-D) as described above with the caveat that the orthogonal evaluation of variants within these regions is much more challenging and our results are certainly less accurate than in the less repetitive regions of the genome. In contrast to the high concordance for deletions in Unique + RM sequences, 30.2% and 59.3% of high confidence deletions from srWGS and lrWGS, respectively, were not shared by the other technology (Figure S6E). The distinct patterns of concordance were more dramatic for insertions: only 17.4% of insertions from lrWGS were overlapped with an srWGS variant, whereas 74.4% of srWGS insertions were captured by lrWGS (Figure S6F). These results highlighted that a major source of added value from lrWGS over srWGS is found in increased SV sensitivity within highly repetitive regions of the genome.

In conclusion, we demonstrate the influence of genomic context on setting expectations for SV detection from srWGS in genomic studies, as well as estimating the anticipated yields of emerging lrWGS technologies. Initial genome-wide surveys have implied highly variable outcomes and limited overall concordance in SV detection between the two technologies; however, in-depth analyses of these variants emphasize that genome organization, variant type, and high type I error rates in SV detection from each technology were the three predominant features driving discordance. After applying *post hoc* filters to correct for the relatively high type I error rates for SV detection from this ensemble srWGS approach optimized for sensitivity and the assembly based lrWGS approach that was optimized with orthogonal data types, we were able to extrapolate the informative biological factors that influenced differences in SV distributions between technologies. The concordance between srWGS and lrWGS was remarkably high (93.8%) for deletions localized to the least-repetitive regions of the genome, while almost all lrWGS-specific deletions were localized to repetitive SD/SR regions.

The value added for long-read assembly to discover new disease associated SVs, or provide resolution to ‘unsolved’ cases in Mendelian genetics, is thus a complex calculus. As we note above, srWGS captures virtually all high-quality deletions derived from lrWGS assembly in the regions of the genome that encompass more than 95% of currently annotated coding sequence in genes with existing evidence for dominant-acting pathogenic mutations from OMIM, so we anticipate that a minority of ‘unsolved’ cases will be explained by cryptic lrWGS SVs from this readily interpretable class of heterozygous deletions in currently known disease-associated genes. However, given that the most highly repetitive regions of the genome have been traditionally inaccessible for genomics studies of disease, it is anticipated that new disease-associated genes and sequences will emerge from these existing blind spots in the human genome. Indeed, germline and somatic repeat expansions and contractions are already well established mechanisms of human disease, particularly neurodegenerative disorders,^42^ and this is an exciting area for future discoveries from lrWGS. As telomere-to-telomere assembly methods continue to mature and eventually reach into centromeres, telomeres, and segmental duplications, the catalogue of disease associated variants will certainly expand beyond what is applied to current clinical interpretation. Similarly, lrWGS was superior for the detection of insertions, irrespective of genomic context, and the near-term value of lrWGS to better delineate coding and noncoding insertions and mobile elements across all genomic contexts is high.

Collectively, we estimate from these analyses that genomic studies and clinical initiatives using srWGS can expect to capture upwards of 10,000 SVs in each human genome, and current large-scale international initiatives are poised to provide exciting new insights into the 90% of the annotated reference genome that encompasses almost all known genic sequence. We also confirm that assembly-based lrWGS methods will access regions of the genome that are intractable to srWGS, and advancements in lrWGS technologies, as well as computational annotation and interpretation tools, will provide significant long-term value in expanding the catalogue of functional variation associated with insertions and mobile elements, as well as variation localized to the most challenging sequence features in the human genome.

## Acknowledgments

Data and analyses were conducted by the Human Genome Structural Variation Consortium (HGSVC). Analyses, data and personnel were supported by the following grants from the National Institutes of Health (NIH): U24HG007497, R01MH115957, R03HD099547, UM1HG008895, R01HD081256, R01HD091797, R01HD096326, R00DE026824, GRFP2017240332, and F31HG010569. C.L. was supported in part by the operational funds from The First Affiliated Hospital of Xi’an Jiaotong University. C.L. is also a distinguished Ewha Womans University Professor supported in part by the Ewha Womans University Research grant of 2019.

## Supplemental Data

There is supplemental information associated with this study, which includes detailed methods, figures and tables. These materials have been provided in a separate document, which will be linked directly from bioRxiv.

## Declaration of Interests

The authors declare no competing interests.

## Web Resources

srWGS data of HGSVC sample, ftp://ftp.1000genomes.ebi.ac.uk/vol1/ftp/data_collections/hgsv_sv_discovery/data/

lrWGS data of HGSVC sample, ftp://ftp.1000genomes.ebi.ac.uk/vol1/ftp/data_collections/hgsv_sv_discovery/working/20160623_chaisson_pacbio_aligns/ VaPoR, https://github.com/mills-lab/vapor

svtk, https://github.com/talkowski-lab/svtk

## Notes

### Competing Interest Statement

The authors have declared no competing interest.

