## Supplemental Material and Methods for "Expectations and blind spots for structural variation detection from short-read alignment and long-read assembly"

### Samples, sequencing, and SV discovery

We utilized three parent-child trios from the 1000 Genomes Project that have been recently analyzed for SVs with both srWGS and lrWGS in the HGSVC.[1] These trios were derived from Han Chinese (CHS), Puerto Rican (PUR) and Yoruban (YRI) Nigerian ancestry groups. The HGSVC generated srWGS and lrWGS data and corresponding SV callsets on these samples, which we used in this study. For srWGS, samples were sequenced with Illumina HiSeq 2500 to ~74.5X coverage per genome, and SVs were discovered using an ensemble approach that integrated 13 independent SV discovery algorithms (WHAMG,[2] LUMPY,[3] DELLY,[4] ForestSV,[5] Manta,[6] Pindel,[7] SVelter,[8] novoBreak,[9] MELT,[10] VariationHunter,[11] dCGH,[12] GenomeSTRiP,[13] and retroCNV[14]). For lrWGS, samples were sequenced with Pacific Biosciences RS II to ~20.0X in the parental genomes and ~39.6X in the child genomes, and SVs were discovered using the integration of two genome assembly-based methods (Phased-SV and MS-PAC[1,15,16]). We used all integrated SV calls as published by the HGSVC, and combined srWGS duplications with insertions for sake of comparisons to lrWGS, which did not distinguish between insertions and duplications.

### SV annotation by repeat content

We defined genomic repeat content including the RM, SD and SR in GRCh38 based on annotations downloaded from the UCSC genome browser (version 2018-08-10).[17] Regions in the RM track that overlapped any SR or SD elements were excluded from RM to avoid conflicting repeat types. Genomic regions falling outside of RM, SR and SD were annotated as "Unique" genomic sequences. We annotated SVs by first allocating their breakpoints to one of the repeat content and assigned each SV to one repeat category by prioritizing SR, followed by SD, RM and then Unique sequences, thus prioritizing annotations with variants that overlapped annotated repeat sequences.

### Statistical test of SV distribution across genomic context

We tested the distribution of SVs across different genomic context against the null hypothesis that SVs are evenly distributed across the genome regardless of the genomic context. Under the null hypothesis 1,056 of the 10,884 SVs from srWGS and 2,408 of the 24,825 SVs from lrWGS were expected in the highly repetitive SD and SR regions that consist 9.7% of the genome, while 5,259 and 17,483 were observed in these regions from srWGS and lrWGS respectively. A chi-square test was performed (Table S1) to test the significance of observation against expectation.

**Comparison of SVs between technologies**

To assess concordance between srWGS and lrWGS, we applied different criteria to SVs based on variant class. We considered deletions to be concordant if over 50% reciprocal overlap of the SV was observed between technologies. Insertions were considered concordant between srWGS and lrWGS if their predicted insertion points were within 100 bp and the lengths of the inserted sequence were within 10 times of each other. As the lrWGS callset did not differentiate duplications from insertions, we also compared lrWGS insertions to srWGS duplications by either 1) requiring >50% of the inserted sequences of lrWGS insertions to be covered by srWGS duplications or 2) requiring >50% reciprocal overlap between the srWGS duplication coordinates and the alignments of assembled lrWGS insertion sequences against the human reference genome (GRCh38) with BLAT(v35).[18] Finally, given that SVs were strictly defined as ≥50bp in the original srWGS and lrWGS SV callsets, we avoided biasing our comparisons near the 50bp size threshold by including small insertions and deletions (indels) defined by both technologies that were between 30-50bp when assessing SV concordance.

**Evaluation and adjudication of SVs**

We designed an *in silico* re-adjudication procedure and applied it to all SVs to reduce the type I error rate of the original SV callsets. We examined orthogonal support from both lrWGS and srWGS data to quantify strength of evidence for each SV. These analyses are described below.

First, to assess raw lrWGS evidence supporting each SV, we applied VaPoR,[19] an algorithm designed to evaluate SV predictions by directly comparing lrWGS sequences with a reference genome through recurrence plots. We executed VaPoR with default settings and considered SVs with a positive genotype score (VaPoR GS >0) as having lrWGS support (Figure 2A, S3A, S5A). In order to maximize validation power, we also applied VaPoR on the parental lrWGS genomes (20.0X) considering SV support in parent as valid. In some regions of the genome

VaPoR is unable to make an evaluation because of nearby sequence homology or low coverage. In this study VaPoR was unable to evaluate 4.1% of srWGS and 5.3% of lrWGS SVs in Unique + RM sequences.

Second, for srWGS data, we focused on three SV signatures: normalized read depth (RD), aberrant paired-end reads (PE), and split reads (SR). RD represents the copy state of a genomic region as relative to expected copy ratio of 1, i.e. RD<1 is related to copy number loss, and RD>1 represents copy number gain. We collected RD, PE and SR evidence per sample using the software package *svtk*.[20] For each SV in Unique + RM sequences, we assessed RD spanning the SV and RD of the 1Kb regions flanking the SV. We next trained an SV classifier using RD values from deletions that were supported by VaPoR as compared to their flanking RD values. For a given RD threshold, we defined the false discovery rate (FDR) as the proportion of flanking regions that had a lower RD threshold and defined the true positive rate (TPR) as the proportion of VaPoR-supported deletions that had a lower RD threshold. We selected a conservative RD cutoff for deletions at 0.35 copy state to keep FDR below 1% (Figure S1) with an understanding this cutoff is optimal for rescuing high-confidence deletions mis-interpreted by VaPoR despite excluding most heterozygous deletions. We applied the same method to duplications and decided a cutoff at 1.60. As expected, RD did not differentiate insertions from their flanking regions (Figure S1), and thus we did not consider RD when filtering insertions.

We similarly determined srWGS PE and SR thresholds to distinguish likely true SVs from false positives. We collected counts of PE reads that were within 100 bp of each breakpoint of an SV and collected SR counts within 50bp from each SV breakpoint. For SVs with more than one breakpoint, we used the minimum PE and SR counts for that SV. Like RD, we designed a classification model as follows. We first generated three training groups: high-confidence homozygous SVs, high-confidence heterozygous SVs, and likely false positive SVs. We defined high-confidence homozygous deletions as those genotyped as homozygous alternative (1/1) by VaPoR, had VaPoR support identified in both parental genomes, and had RD of 0. High-confidence heterozygous deletions were genotyped as heterozygous (0/1) by VaPoR, had VaPoR support in only one parental genome, and had RD between 0.45 and 0.5. Likely false positive deletions did not have VaPoR support in any genomes in the trio, had RD >1, and were labeled as *de novo* in the original callset for SVs from srWGS. Duplications with RD >1.6 that were genotyped as 1/1 by VaPoR and had VaPoR support in both parental genomes were considered high-confidence homozygous duplications. Those that displayed RD>1.6, were genotyped as 0/1 by VaPoR and had VaPoR support in only one parental genome were considered high-confidence heterozygous duplications, and those that lack VaPoR support in all trio genomes, had RD <1.6 and were labeled as *de novo* in the original srWGS callset were considered likely false positives. For insertions, we relied solely on VaPoR results to define srWGS training sets. We defined high-confidence homozygous insertions as those genotyped as homozygous by VaPoR and had support in both parental genomes. We defined high-confidence heterozygous insertions as those genotyped as heterozygous by VaPoR and had VaPoR support in only one parental genome. Finally, we defined likely false-positive insertions as those without VaPoR support in any genomes in the trio (Figure S2).

After identifying the SV subsets defined above for srWGS PE/SR classifier training, we varied thresholds for PE and SR to seek optimal values for each type of SVs by restricting the FDR to <1%, defined as the proportion of likely false-positive SVs that have more PE and SR support than the selected threshold, while maximizing the TPR, defined as proportion of high-confidence homozygous and heterozygous SVs that have higher PE and SR support. As explicated in Table S2, we selected PE and SR thresholds at 15 and 0, respectively, for deletions, resulting in FPR of 0.95% and TPR of 92.59% for homozygous and 97.52% for heterozygous deletions observed. Comparable results are displayed for duplications and insertions in Table S2.

### Supplemental Figures and Legends

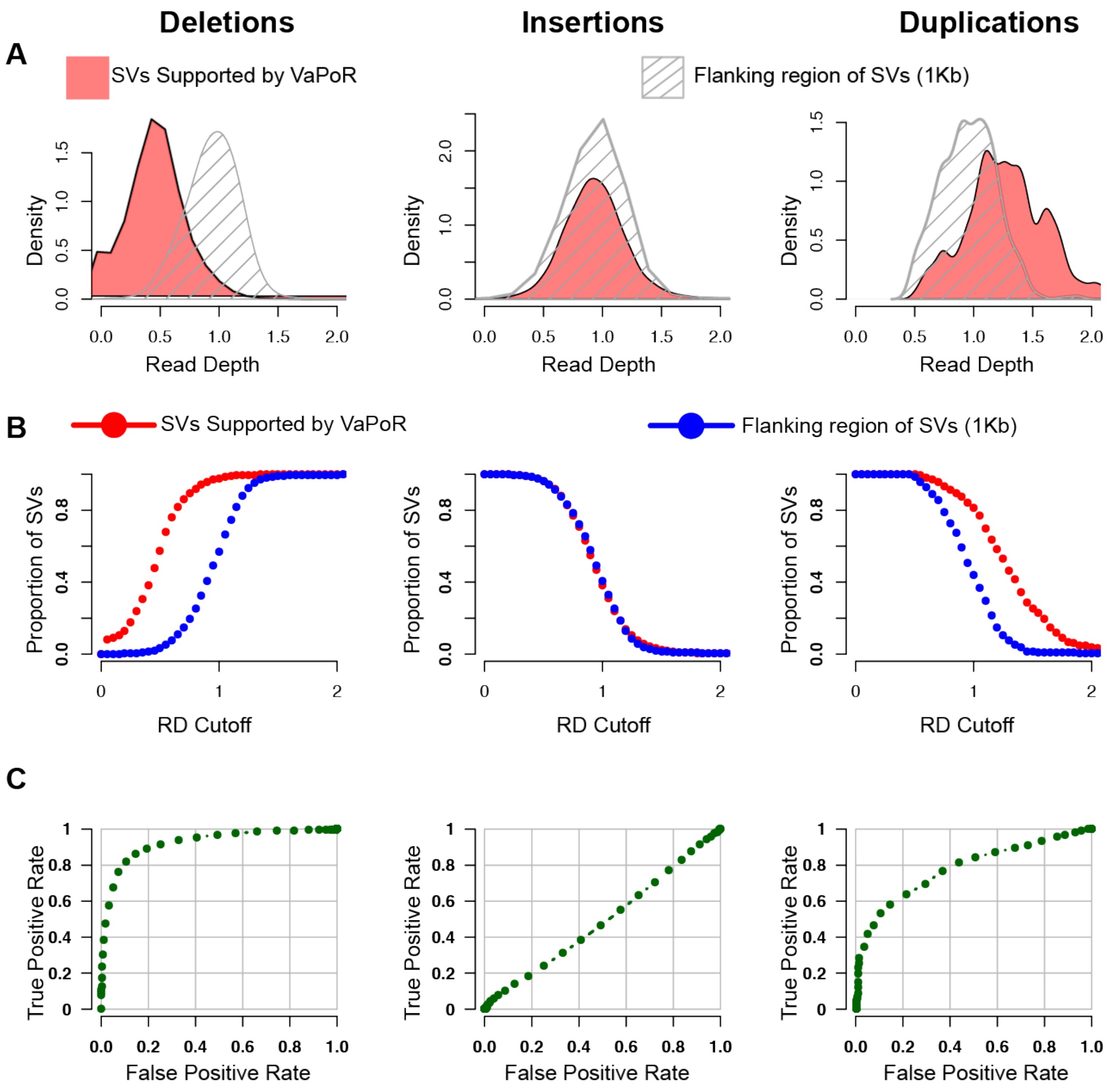


**Figure S1. Distribution of normalized read depth of srWGS for SVs supported by long reads.**

(A) Distribution of normalized read depth (RD) for deletions (left), insertions (middle) and duplications (right) that were supported by VaPoR (red) and the 1Kb flanking regions of these SVs (grey).

(B) Proportion of deletions (left), insertions (middle) and duplications (right) that passed the different cutoffs of RD (true positive rate, red), and the proportion of the 1Kb flanking regions of these SVs that passed the cutoffs (false positive rate, blue).

(C) Receiver operating characteristic (ROC) of RD for deletions (left), insertions (middle) and duplications (right). True positive and false positive rates are as defined in (B).

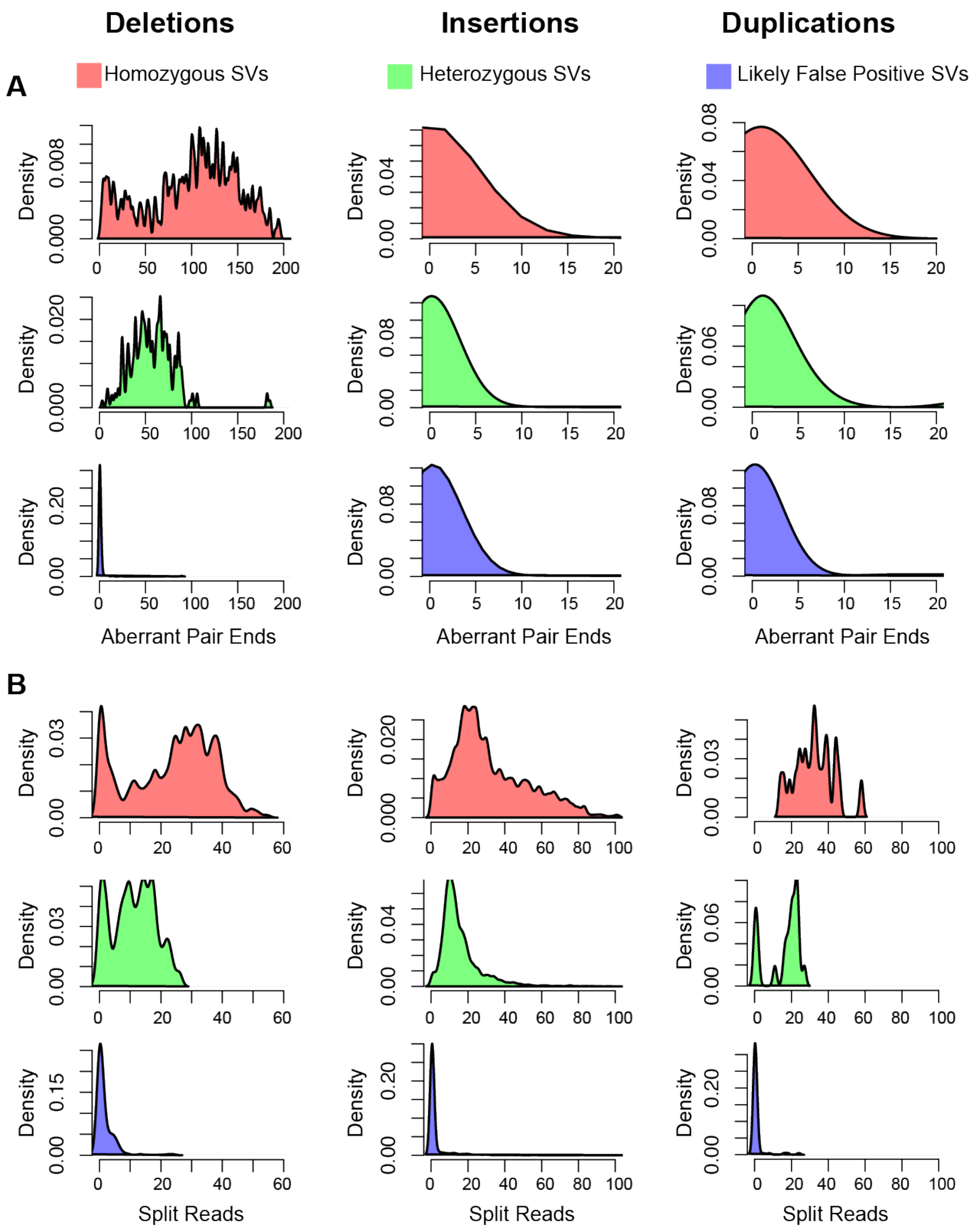


**Figure S2. Distribution of aberrant paired-end reads (PE) and split reads (SR) from srWGS across high confidence homozygous SVs, heterozygous SVs and likely false positive SVs.**

Distributions of (A) PE and (B) SR metrics for homozygous (red), heterozygous (green) and false positive (blue) SVs for deletions (left), insertions (middle), and duplications (right).

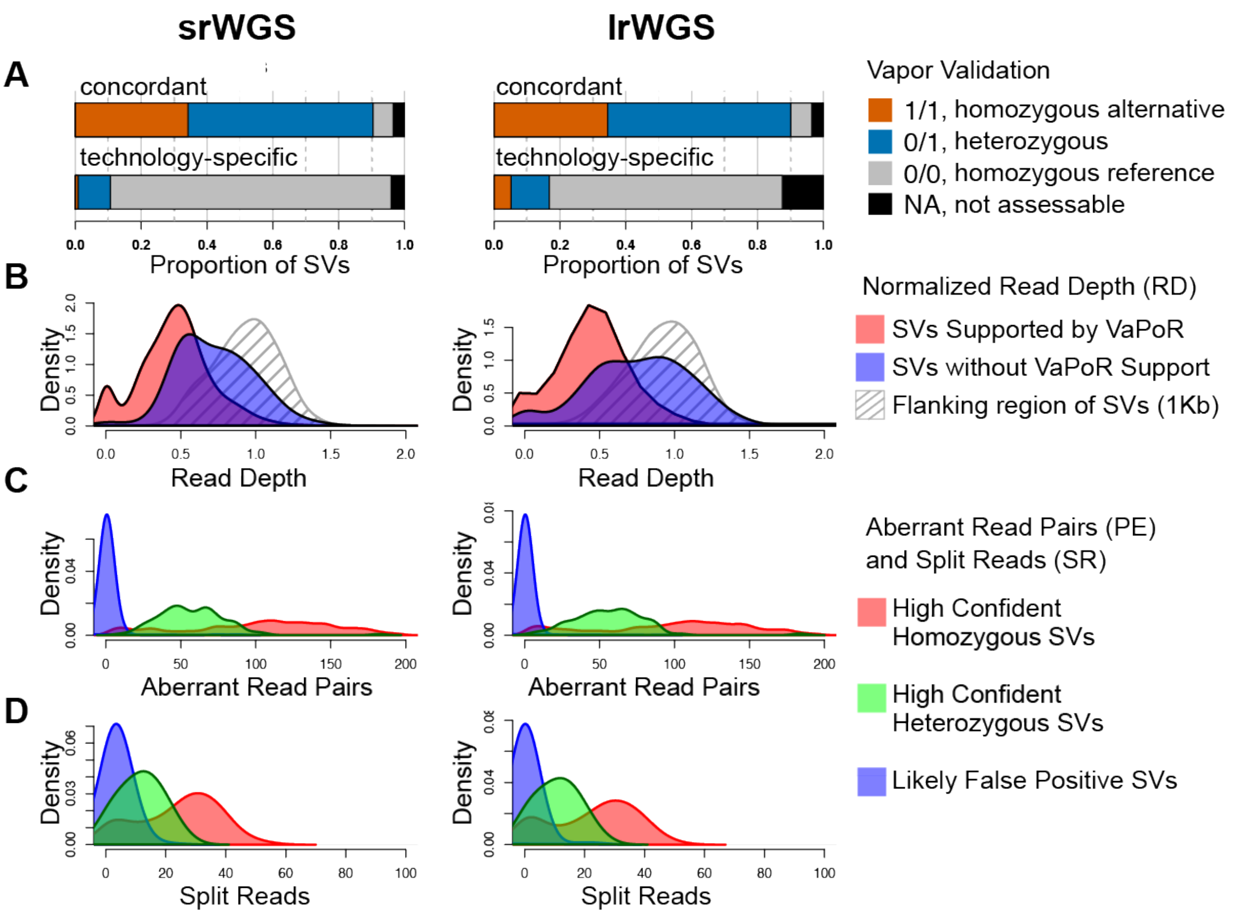


**Figure S3. Error correction methods for deletions from srWGS and lrWGS based on read-level alignment signatures.**

(A) *In silico* evaluation results from VaPoR on deletions from srWGS (left) and lrWGS (right).

(B) Distribution of normalized read depth (RD) of srWGS across deletions that have support from VaPoR (red), deletions that do not have support from VaPoR (blue), and the 1Kb regions that flank each deletion (grey).

(C-D) Distribution of (C) aberrant srWGS read pairs and (D) split reads from srWGS alignments around deletion breakpoints that were either homozygous (red), heterozygous (green) or likely false positives (blue).

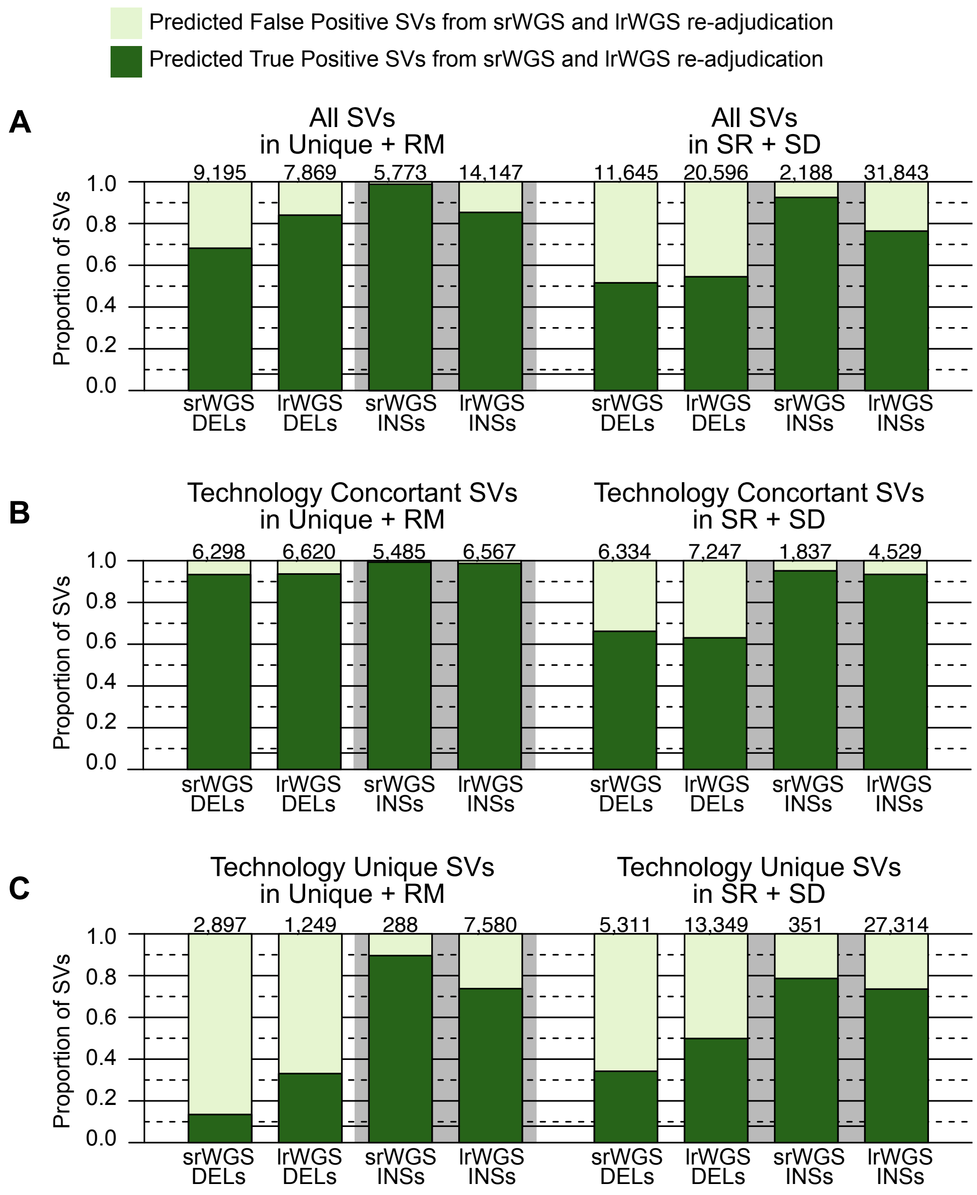


**Figure S4. Proportion of SVs supported by the *in silico* refinement procedure.**

(A) Proportion of deletions (DELs) and insertions (INSs) that were supported by the *in silico* refinement procedure.

(B) Proportion of SVs concordant between srWGS and lrWGS that were supported by the *in silico* refinement procedure.

(C) Proportion of SVs uniquely discovered by srWGS or lrWGS that were supported by the *in silico* refinement procedure.

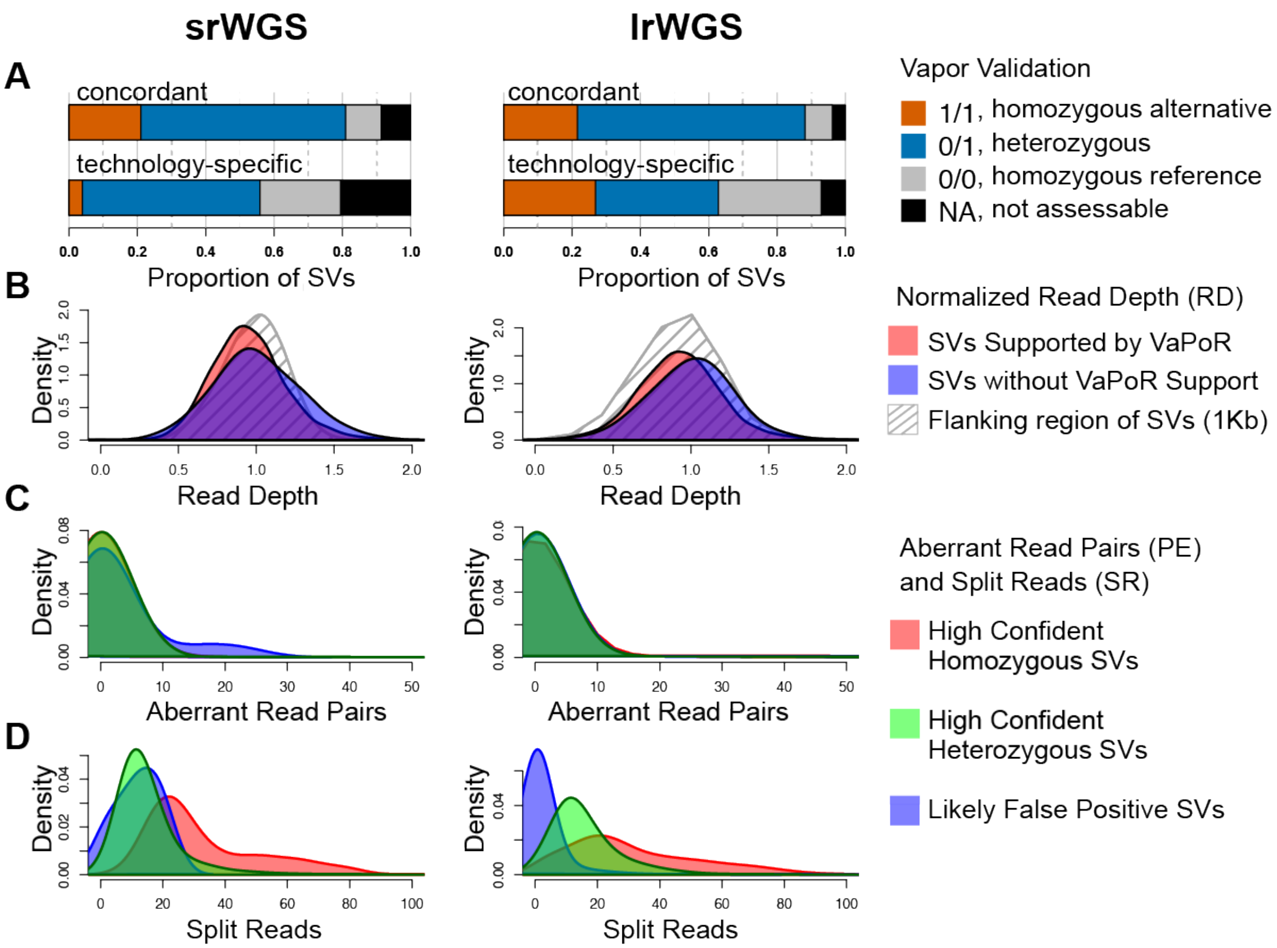


**Figure S5. Error correction methods for insertions from srWGS and lrWGS based on read-level alignment signatures.**

(A) *In silico* evaluation results from VaPoR on insertions from srWGS (left) and lrWGS (right).

(B) Distribution of normalized read depth (RD) of srWGS across insertions that have support from VaPoR (red), insertions that do not have support from VaPoR (blue), and the 1Kb regions that flank each insertion (grey).

(C-D) Distribution of (C) aberrant srWGS read pairs and (D) split reads from srWGS alignments around insertion breakpoints that were either homozygous (red), heterozygous (green) or likely false positives (blue).

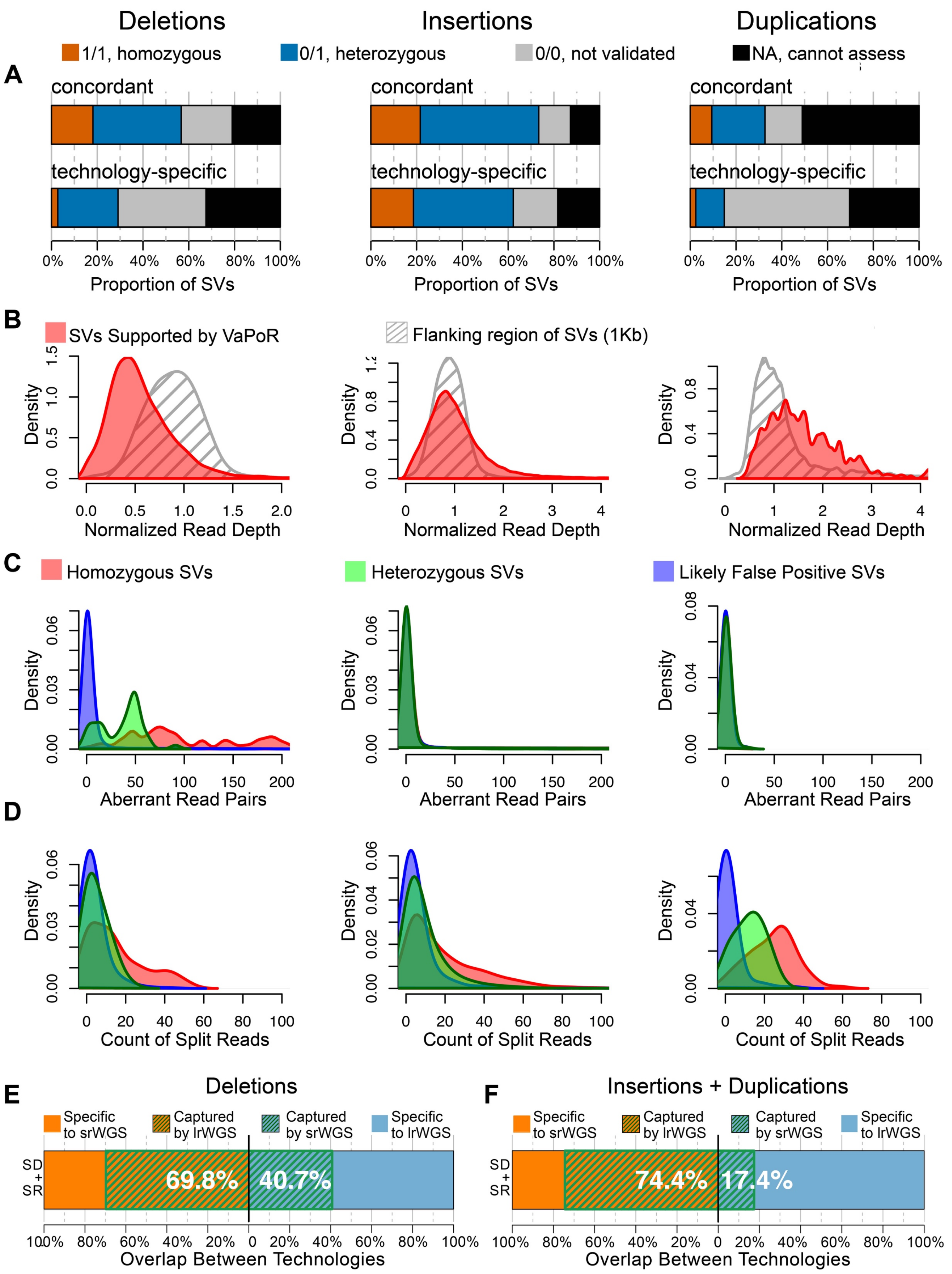

Deletions
Insertions
Duplications
1/1, homozygous
0/1, heterozygous
0/0, not validated
NA, cannot assess
A
concordant
technology-specific
0%
20%
40%
60%
80%
100%
Proportion of SVs
B
SVs Supported by VaPoR
Flanking region of SVs (1Kb)
Density
Normalized Read Depth
C
Homozygous SVs
Heterozygous SVs
Likely False Positive SVs
Aberrant Read Pairs
D
Count of Split Reads
E
Deletions
Specific to srWGS
Captured by lrWGS
Captured by srWGS
Specific to lrWGS
SD + SR
69.8%
40.7%
Overlap Between Technologies
F
Insertions + Duplications
74.4%
17.4%

**Figure S6. Error correction method for SVs in SD+SR region and the updated concordance.**

(A) *In silico* evaluation results from VaPoR on deletions (left), insertions (middle) and duplications (right). Deletions and insertions were reported in both srWGS and lrWGS callsets; duplications were only reported in the srWGS callset.

(B) Distribution of normalized read depth of srWGS across deletions (left), insertions (middle) and duplications (right) that were supported by VaPoR (red), and the 1Kb genomic regions that flank each SV (grey).

(C-D) Distribution of (C) aberrant srWGS read pairs and (D) split reads around deletions (left), insertions (middle) and duplications (right) that were either homozygous (red), heterozygous (green) or false positives (blue). The homozygous, heterozygous and likely false positive SV sets were selected using the criteria described in supplemental methods.

(E-F) Concordance of (E) deletions and (F) insertions and duplications in SD+SR sequences that were supported by the *in silico* SV refinement procedure. Percentages represent the fraction of total variants shared between srWGS and lrWGS.

### Supplemental Tables

**Table S1. Expected and observed counts of SVs located within SD + SR, and in Unique + RM sequences.**

| | srWGS | | lrWGS | |
|---|---|---|---|---|
| | SR + SD | Unique + RM | SR + SD | Unique + RM |
| Expected | 1056 | 9828 | 2408 | 22417 |
| Observed | 5259 | 5625 | 17483 | 7342 |

**Table S2. PE and SR cutoffs selected to discriminate the quality of SVs from lrWGS and srWGS callsets.**

| | Selected Thresholds | | SVs Passing Thresholds in each Training Set (%) | | |
|---|---|---|---|---|---|
| Type of SVs | PE | SR | Homozygous | Heterozygous | Predicted Type I Errors |
| DEL | 15 | 0 | 92.59% | 97.52% | 0.95% |
| DUP | 0 | 19 | 73.81% | 57.14% | 1.22% |
| INS | 0 | 30 | 42.87% | 8.25% | 0.94% |

association studies and its implications for autism spectrum disorder. Nat. Genet. *50*, 727–736.
